# Generation of an inducible destabilized-domain Cre mouse line to target disease associated microglia

**DOI:** 10.1101/2024.09.18.613773

**Authors:** Caden M. Henningfield, Nellie Kwang, Kate Inman Tsourmas, Jonathan Neumann, Shimako Kawauchi, Vivek Swarup, Grant R. MacGregor, Kim N. Green

## Abstract

The function of microglia during progression of Alzheimer’s disease (AD) can be investigated using mouse models that enable genetic manipulation of microglial subpopulations in a temporal manner. We developed a mouse strain that expresses destabilized-domain Cre recombinase (DD-Cre) from the *Cst7* locus (*Cst7*^DD-Cre^) and tested this in 5xFAD amyloidogenic, Ai14 tdTomato cre-reporter line mice. Dietary administration of trimethoprim to induce DD-Cre activity produces long-term labeling in disease associated microglia (DAM) without evidence of leakiness, with tdTomato-expression restricted to cells surrounding plaques. Using this model, we found that DAMs are a subset of plaque-associated microglia (PAMs) and their transition to DAM increases with age and disease stage. Spatial transcriptomic analysis revealed that tdTomato+ cells show higher expression of disease and inflammatory genes compared to other microglial populations, including non-labeled PAMs. This model should allow inducible cre-loxP targeting of DAMs, without leakiness.

**Highlights:**

- We developed a new mouse strain which specifically enables recombination of loxP sites in disease associated microglia (DAMs) and can be used to manipulate DAM-gene expression.
- DAMs represent a subset of plaque associated microglia (PAMs), and DAM expression increases with disease progression.
- Spatial transcriptomic analyses reveal that DAMs have higher expression of disease and inflammatory genes compared to other PAMs.

## Introduction

Alzheimer’s disease (AD), the most common human neurodegenerative disease, is characterized by the appearance of amyloid-beta (Aβ) plaques and neurofibrillary Tau tangles (NFT). In AD, microglia, the resident immune cells of the brain, cluster around Aβ plaques in the brain parenchyma in what is theorized as an attempt to control and restrict plaques and limit dystrophic neurites ^1–3^. Numerous additional roles for microglia in the pathogenesis of AD have been postulated including in plaque seeding and formation ^4^, synaptic, neuronal, and perineuronal net loss ^4–7^, facilitation of tau pathology and spreading ^8,9^, and downstream neurodegeneration ^10,11^. Consistent with a role for microglial function in AD, genome wide association studies (GWAS) have identified single nucleotide polymorphisms (SNPs) in genes whose expression is highly enriched in microglia and which confer a heightened risk of developing AD such as *TREM2*, *SPI1*, *PLCG2*, *ABI3*, amongst others ^12–16^.

Microglia are highly plastic and will rapidly remodel their gene expression and physical properties upon sensing changes in their local environment. In the AD brain this can occur in response to Aβ, plaques, tau-laden neurons, or cell death, and results in dramatic changes in microglial phenotypes. The assumption is that these changes underlie facets of the degeneration process and understanding each change may provide valuable insight into the disease process. These changes have been illustrated by single-cell RNA sequencing which has provided unprecedented insight into the heterogeneity of microglia, identifying transcriptionally distinct microglial sub-populations, including but not limited to DAMs ^3^, injury responsive microglia (IRM) ^17^, axon tract associated microglia (ATM) ^17–20^, and white matter associated microglia (WAM) ^21^. Of particular interest, DAMs are characterized by increased expression of inflammatory genes such as *Trem2, Cst7, Lpl, Axl, Clec7a, Itgax, Ccl6,* and others, and likely represent plaque associated microglia (PAMs) in the AD brain ^3^. However, the exact relationship between DAMs and PAMs is not yet fully understood and whether DAMs represent all, or just a subset of PAMs is unclear. These studies suggest that microglia have distinct signaling pathways and functions based on their location within the brain and their association with disease/injury.^3,17,21^

Investigating gene function in DAMs is of interest to both basic research as well as screening for targets for therapeutic intervention of AD. Current approaches to manipulate gene expression in microglia target all populations making it difficult to analyze the function of a particular gene during the response of the microglia to pathology versus its involvement in homeostatic functions. For example, TREM2 mediates the microglial response to plaques, as shown by its knockout in models of amyloidosis ^3,22,23^, but loss of TREM2 function also causes perturbations in homeostatic microglial function leading to the neurodegenerative disease Nasu Hakola ^24,25^. With this in mind, we sought to develop tools which could be used to modulate gene function in DAM versus homeostatic microglia.

To that end, we developed a novel, inducible Cre recombinase mouse line which expresses destabilized-domain Cre (DD-Cre) from the *Cst7* locus, whose expression is highly upregulated in nascent DAM, allowing us to target these disease-responsive microglia with temporal specificity. We show that our model specifically targets DAMs, and using spatial transcriptomics we demonstrate that DAMs represent a subset of the entire PAM population, illustrating the importance of determining the exact roles that diverse microglial subsets have in AD.

## Methods

### Animals

All animal experiments performed in this study were approved by the UC Irvine Institutional Animal Care and Use Committee (IACUC) and were compliant with ethical regulations for animal research and testing. The 5xFAD mouse strain ((B6.CgTg(APPSwFlLon,PSEN1*M146L*L286V)6799Vas/Mmjax, Jackson Lab Stock # 34848, MMRRC)) has been described in detail ^26^. All mice used in this study were on a congenic or co-isogenic C57BL/6J background.

### Environmental conditions

Animals were housed in autoclaved individual ventilated cages (SuperMouse 750, Lab Products, Seaford, DE) containing autoclaved corncob bedding (Envigo 7092BK 1/8” Teklad, Placentia, CA) and two autoclaved 2” square cotton nestlets (Ancare, Bellmore, NY) plus a LifeSpan multi-level environmental enrichment platform. Tap water (acidified to pH2.5-3.0 with HCl then autoclaved) and food (LabDiet Mouse Irr 6F; LabDiet, St. Louis, MO) were provided *ad libitum*. Cages were changed every 2 weeks with a maximum of 5 adult animals per cage. Room temperature was maintained at 72 ± 2°F, with ambient room humidity (average 40-60% RH, range 10 - 70%). Light cycle was 14h light / 10h dark, lights on at 06.30h and off at 20.30h.

### Generation of mice with a *Cst7-DD-Cre* allele

CRISPR/Cas9 was used to insert a 1599 bp DNA sequence encoding ecDHFR-Cre-P2A immediately downstream of the translation initiation codon in exon 1 of *Cst7*. Alt-R Crispr RNA (TMF1132 – AAGCAGAATGGCCAGCCACA) and tracrRNA plus CAS9 protein (HiFi Cas9 nuclease V3, Integrated DNA Technologies (IDT), Coralville, IA) as a ribonucleoprotein (RNP) was microinjected into C57BL/6J zygotes (Jackson Lab Stock # 000664) along with a 1805 base ssODN sequence (TMF1130 – Supplementary File 1) to introduce the DD-Cre-P2A coding sequence into exon 1 of the *Cst7* locus immediately downstream of the initiator ATG sequence. G0 founder animals containing the desired DNA sequence changes were backcrossed with C57BL/6J mice and N1 heterozygous mice were sequenced to determine the variant allele. N1 heterozygous mice were backcrossed to produce N2 heterozygotes, which were used to generate animals for subsequent analysis.

#### Genotyping of mouse alleles

Oligonucleotides for PCR-based genotyping were purchased from IDT. The 5xFAD transgene was identified using PS1 Forward 5′ - AAT AGA GAA CGG CAG GAG CA – 3′ and PS1 Reverse 5′ - GCC ATG AGG GCA CTA ATC AT – 3′. The Cst7^DD-cre^ and WT alleles, were identified using the following primers: WT *Cst7* F 5’ – CCCAAGTCCTGAAGATGAAGCG - 3’, WT *Cst7* R 5’ – CCACCGCCTGATCTATGGTG – 3’, and ecDHFR R 5’ – GCATAGCGTTTTCCATCCCG – 3’ with a 206 bp product for the *Cst7*^DD-cre^ allele and 344 bp for the WT *Cst7* allele. The Ai14^tdTomato^ allele^27^ was identified using WT Forward 5’ – AAGGGAGCTGCAGTGGAGTA – 3’, WT Reverse 5’ – CCGAAAATCTGTGGGAAGTC – 3’, Mutant Reverse 5’ – GGCATTAAAGCAGCGTATCC - 3’, and Mutant Forward 5’ – CTGTTCCTGTACGGCATGG – 3’ with 196bp product for the Ai14^tdTomato^ allele and 297 bp for the WT allele.

### Animal Treatments

Two-and-a-half-month-old and 9.5-month-old 5xFAD hemizygous, double heterozygous *Cst7*^DD-cre,^ Ai14^tdTomato^ mice were treated with trimethoprim (TMP; Sigma Aldrich, T0667) at 0.8mg/mL in their drinking water for 1.5 months. At the end of treatments, mice were euthanized via CO_2_ inhalation and transcardially perfused with 1X phosphate buffered saline (PBS; Sigma Aldrich, P4417). For all studies, brains were removed, and hemispheres separated along the midline. Brain halves were either flash frozen for subsequent biochemical analysis or drop-fixed in 4% paraformaldehyde in 1X PBS (PFA; Thermo Fisher Scientific, J19943.K2) for subsequent immunohistochemical analysis. Half brains collected into 4% PFA for 48 hr were then transferred to a 30% sucrose solution with 0.02% sodium azide for another 48-72 hr at 4 deg C. Fixed half brains were sliced at 40 µm using a Leica SM2000 R freezing microtome and sliced tissue was stored in 30% glycerol, 30% ethylene glycol, and 1xPBS solution.

#### Cuprizone (CPZ) treatment

Ten-month-old *Cst7*^DD-Cre^/Ai14^tdTomato^ double heterozygous mice were fed 0.3% cuprizone chow (Envigo, Indianapolis, IN) and 0.8mg/mL TMP through drinking water for 6 weeks. Weights of individual mice and chow consumption in each cage were recorded and chow was changed every 7 days to monitor expected weight loss as well as ensuring freshness of cuprizone chow. Brains were collected and fixed in 4% PFA for 48 hr followed by cryoprotection by 30% sucrose 48 hr, all at 4 °C before sectioning.

### Immunohistochemistry

Indirect fluorescence immunolabeling was performed as described ^28,29^. Primary antibodies used were anti-ionized calcium-binding adapter molecule 1 (IBA1; 1:1000 dilution; 019–19741; Wako, Osaka, Japan), anti-CD11c (1:200, 50-112-2633; eBioscience), and anti-CST7 (courtesy of Dr. Collin Watts, University of Dundee, UK). Thioflavin-S (Thio-S; 1892; Sigma-Aldrich) and Amylo-Glo (TR-300-AG; Biosensis, Thebarton, South Australia, AU) were used to visualize Aβ plaques according to manufacturer’s instructions. For Amylo-Glo staining, tissue sections were washed in 70% ethanol for 5 min followed by a 2 min wash in distilled water (all washes were at room temperature unless stated). Sections were then reacted with 1% Amylo-Glo for 10 min then washed with 0.9% 1X PBS for 5 min then distilled water for 15 sec. For Thio-S staining, tissue sections were incubated in 0.5% Thio-S diluted in 50% ethanol for 10 min. Sections were then washed twice for 5 min each in 50% ethanol, then one 10 min wash in 1X PBS before continuing with fluorescent immunolabelling. Sections were then briefly rinsed in 1X PBS and immersed in normal goat or donkey serum blocking solution (5% normal serum with 0.2% Triton-X100 in 1X PBS) for 60 min. Sections were incubated overnight in primary antibody at the dilutions described above in normal serum blocking solution at 4 deg C. The next day tissue sections were washed thrice in 1X PBS for10 min each before being placed in appropriate secondary antibody in normal serum blocking solution (1:200 for all species and wavelengths; Invitrogen) for 60 min. Tissue sections were then washed thrice for 10 min in 1X PBS before tissue was mounted and coverslipped. High resolution fluorescent images were obtained using a Leica TCS SPE-II confocal microscope and LAS-X software. To capture fluorescence images to be assembled into whole brain stitches, automated slide scanning was performed using a Zeiss AxioScan.Z1 slidescanner, equipped with an Orca Flash 4.0 v3 sCMOS camera (Hamamatsu, Bridgewater, NJ) and Zen AxioScan 2.3 software.

### Data Analysis and Statistics

Both male and female mice were used in all statistical analyses. Statistical analysis was accomplished using Prism GraphPad (v9.0.0). To compare two groups, the unpaired Student’s t-test was used. To compare two or more groups at a single timepoint or in a single brain region, one-way ANOVA with Tukey’s multiple comparison correction was performed. To compare two or more groups at either different timepoints or brain regions, with multiple treatments/genotypes, two-way ANOVA with Tukey’s multiple comparison correction was used. For all analyses, statistical significance was accepted at p < 0.05. and significance expressed as follows: *p < 0.05, **p< 0.01, ***p < 0.001.

### Single-cell Spatial Transcriptomics

Isopentane fresh-frozen brain hemispheres were embedded in optimal cutting temperature (OCT) compound (Fisher, 23-730-571), and 10 um thick coronal sections prepared using a cryostat (Leica CM1950). Six hemibrains were mounted directly onto a VWR Superfrost Plus microscope slide (Avantor, 48311-703) and stored at -80°C until fixation. 5xFAD hemizygous, *Cst7*^DD-Cre^, Ai14^tdTomato^ double heterozygous (n=3), and *Cst7*^DD-Cre^/Ai14^tdTomato^ double heterozygous (n=1) mice treated with TMP, and *Cst7*^DD-Cre^/Ai14^tdTomato^ double heterozygous mice treated with cuprizone (CPZ) and TMP (n=2) were used for spatial transcriptomics. Tissues were processed according to the Nanostring CosMx fresh-frozen slide preparation manual for RNA and protein assays.

### Tissue preparation for spatial transcriptomics

Slides were fixed in 10% neutral buffered formalin (NBF; EMS Diasum, 15740-04) for 2 hr at 4°C, followed by three 5 min washes in 1X PBS and dehydration with one 5 min wash in each of 50%, 70% and 100% ethanol. Antigen retrieval was performed at 100°C for 15 min with CosMx Target Retrieval Solution followed by tissue permeabilization for 30 min and fiducial application. After post-fixation with 10% NBF and NHS-Acetate (Fisher, 26777), *in situ* hybridization was performed using the 1000-plex Mouse Neuro RNA panel and rRNA segmentation marker overnight for 16-18 hr in a hybridization oven at 37°C. Two stringent washes with 50% formamide and 4X Saline-Sodium citrate (SSC) solution at 37°C for 30 min each were done prior to nuclear DAPI stain (2.5%) for 15 min. Tissues were incubated in GFAP (4.0%) and histone (4.0%) cell segmentation markers for 1 hr before flow cells were assembled according to the CosMx manual. Once assembled, flow cells were loaded into the CosMx machine with roughly 300 field-of-views (FOVs) selected per slide, capturing the cortex and hippocampus. Slides were imaged for 7 days before the data were uploaded to the Nanostring AtoMx platform for further analysis and visualization. Pre-processed data was then exported as a Seurat object for further analysis in R v4.3.1.

### Spatial transcriptomics data analysis

Spatial transcriptomics datasets were filtered using the AtoMx RNA Quality Control module to flag poorly performing probes, cells, FOVs, and target genes. Datasets were then normalized and scaled using Seurat SCTransform to account for differences in library size across cell types^30,31^. Principal component analysis (PCA) and uniform manifold approximation and projection (UMAP) analysis were performed to reduce dimensionality for downstream analysis. Unsupervised clustering at 1.0 resolution yielded 38 clusters for the dataset. Clusters were manually annotated based on gene expression and spatial location. Differential gene expression analysis per cell type between genotypes was performed using MAST to calculate the average difference, defined as the difference in average expression levels between two conditions^32^. Differential upregulation (DU) and differential downregulation (DD) scores were calculated by summing the product of the negative log10(padj) and the average difference for each statistically significant gene (padj < 0.05) with an absolute average difference greater than 0.3. Microglia were then subsetted and further subclustered for deeper analysis. Data visualizations were generated using ggplot2^33^.

## Results

### Generation of a novel destabilized Cre (DD-Cre) mouse line to target DAMs

To develop a mouse strain that can be used to selectively target and modulate gene expression in DAMs via expression of cre recombinase we screened for genes that were upregulated by microglia around plaques, but which were not expressed under homeostatic conditions. We used data from microglia-depleted 5xFAD mice via chronic administration of CSF1R inhibitor^4^ to identify 10 candidate genes that displayed increased RNA levels in 5xFAD mice, but nearly abolished levels after microglial depletion: *Ccl6, Clec7a, Cst7, Ctsd, Ctsz, C1qa, Hexb, Siglech, Sip1,* and *Itgax*^4^. From this list we screened for genes with undetectable expression in the wild-type brain and found *Cst7* to be a suitable candidate as it showed high expression in 5xFAD mice, but severely reduced levels when microglia were eliminated, and undetectable expression in the WT brain^4^. Having identified *Cst7* as an appropriate driver, we wanted to develop a model in which DAM’s could be targeted inducibly throughout the animal’s lifespan and without utilization of tamoxifen, which is potentially toxic, poor at crossing the BBB, and has effects on macrophage populations^34–37^. An attractive alternative to tamoxifen-inducible cre recombinase relies on the ability of the antibiotic trimethoprim (TMP) to promote stability of a mutant destabilized domain from bacterial dihydrofolate reductase (ecDHFR) fused to cre recombinase, known as DD-Cre^37^. In the absence of TMP, DD-Cre is normally degraded through the proteasomal pathway. However, in the presence of TMP, the DD-Cre fusion protein is stabilized, and its decay is blocked allowing it to mediate recombination at l*oxP* sites. TMP is safer for mice compared to tamoxifen and it can also be dissolved into the animal’s drinking water allowing for long-term treatment, unlike tamoxifen which can only safely be administered short-term.

Cell-type specific driver lines of mice in which Cre recombinase has been knocked into a locus often produces haploinsufficiency of the target gene, which can confound experimental analysis. To overcome this potential issue, we used CRISPR/Cas9 to insert a DD-Cre coding sequence at the ATG translation initiation codon of the mouse *Cst7* locus followed by a P2A ribosomal skip coding sequence (Supplementary File 1). In this *Cst7*^DD-Cre^ allele (Fig. 1A), both DD-CRE and CST7 proteins are expressed under control of the *Cst7 cis*-regulatory elements. G0 founder mice were screened for the presence of the DD-Cre sequence at the *Cst7* locus. Founder animals were bred with C57BL/6J mice and N1 heterozygous animals with the desired sequence modification were identified. Heterozygous *Cst7*^DD-Cre^ N1 generation mice were crossed again with wildtype C57BL/6J mice to segregate any potential unlinked mutations. N2 heterozygous *Cst7*^DD-Cre^ mice were used for subsequent studies.

**Figure 1:**
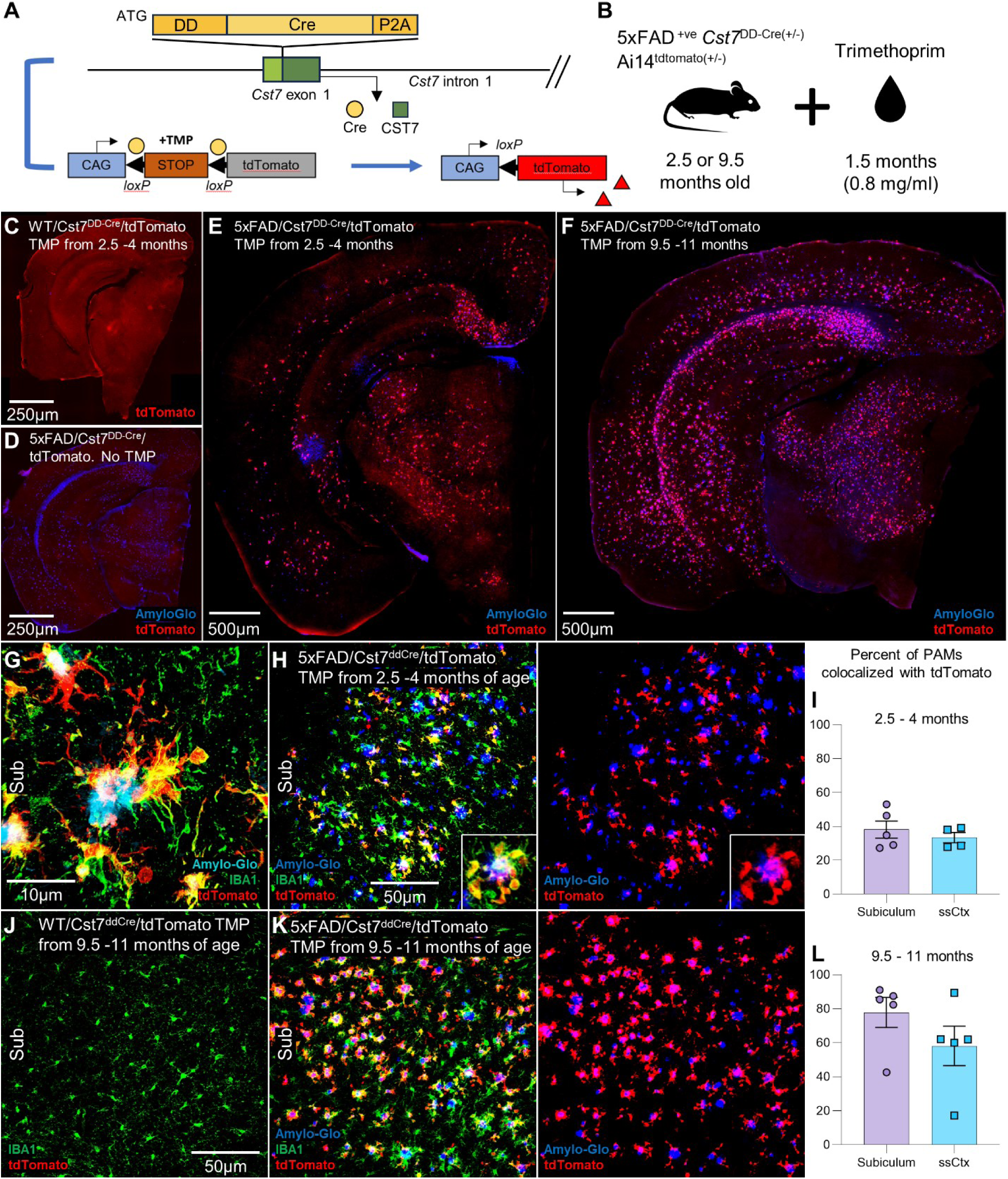
Novel *Cst7*^DD-Cre^ mouse line specifically targets plaque associated microglia (PAMs). (A) Diagram showing location of DD-Cre-P2A coding sequence inserted immediately upstream of the initiator AUG codon in exon 1 of *Cst7*, plus genetic paradigm used to assess lineage tracing of DD-Cre activity. When trimethoprim (TMP) is administered, the stabilized DD-Cre recombines *loxP* sites flanking the STOP cassette preceding the tdTomato genetic sequence. tdTomato is subsequently expressed only in cells that have expressed *Cst7*. (B) Schematic showing experimental paradigm. Two and a half and 9.5 month-old 5xFAD/*Cst7*^DD-Cre^/Ai14^tdTomato^ mice were treated with TMP in drinking water for 1.5 months before sacrifice. (C-F) Representative images of *Cst7*^DD-Cre^/Ai14^tdTomato^ negative control mice treated with TMP showing weak autofluorescence (C), and 5xFAD/*Cst7*^DD-Cre^/Ai14^tdTomato^ experimental mice not treated with TMP (D) show no expression of tdTomato. By contrast, 4-month (E) and 11-month (F) old 5xFAD/*Cst7*^DD-Cre^/Ai14^tdTomato^ mice treated with TMP show multiple cells expressing tdTomato. (G) Example of a 5xFAD/*Cst7*^DD-Cre^/Ai14^tdTomato^ mouse treated with TMP showing tdTomato colocalization with microglial marker, IBA1, and plaque stain, Amylo-Glo. (H-I) Example of subiculum in a 4-month-old 5xFAD/*Cst7*^DD-Cre^/Ai14^tdTomato^ mouse showing cells expressing tdTomato in proximity to plaques and microglial colocalization (H). Quantification of the average percentage of PAMs that colocalize with tdTomato is 38% and 33% for the subiculum and somatosensory cortex, respectively (I). (J) Example of absence of tdTomato expression in *Cst7*^DD-Cre^/Ai14^tdTomato^. control mouse treated with TMP (K-L) Example of subiculum in an 11-month-old 5xFAD/*Cst7*^DD-Cre^/Ai14^tdTomato^ mouse illustrating tdTomato proximity to plaques and microglial colocalization (K). Quantification of the percentage of PAMs that colocalize with tdTomato is an average of 78% and 58% for the subiculum and somatosensory cortex, respectively (L).

### *Cst7*^DD-Cre^ specifically targets DAMs in 5xFAD transgenic mice

To investigate whether the *Cst7*^DD-Cre^ line specifically and inducibly targets cre activity to *Cst7* expressing cells, we bred these mice to the 5xFAD (Tg6799) mouse model of amyloidosis^26^ and subsequently bred their offspring with a loxP-stop-loxP tdTomato reporter line knocked into the Gt(ROSA)26Sor (ROSA26) locus (Ai14^tdTomato^; Fig. 1A) to generate 5xFAD hemizygous, *Cst7*^DD-Cre^ heterozygous, Ai14 tdTomato heterozygous (5xFAD/*Cst7*^DD-Cre^/Ai14^tdTomato^) experimental and *Cst7*^DD-Cre^ heterozygous, Ai14 tdTomato heterozygous (*Cst7*^DD-Cre^/Ai14^tdTomato^) control mice. In the absence of cre activity, the loxP-flanked stop (LSL) cassette in the Ai14 tdTomato allele prevents transcription of the downstream tdTomato sequence, as the LSL cassette mediates transcriptional termination via three tandem SV40 polyA signals. However, upon stabilization of the DD-Cre protein via TMP, DD-Cre mediates removal of the LSL cassette enabling expression of tdTomato via the CAG promoter (Fig. 1A). To analyze early and late responses to plaques, we treated 2.5- and 9.5-month-old 5xFAD/Cst7 ^DD-Cre^/Ai14^tdTomato^ mice with TMP for 1.5 months at a concentration of 0.8mg/ml in their drinking water at which point mice were sacrificed at 4- and 11-months of age, respectively (Fig. 1B).

As predicted, *Cst7*^DD-Cre^/Ai14^tdTomato^ (non-5xFAD) control mice treated with TMP show only background weak autofluorescence (i.e. no tdTomato expression) in the brain (Fig. 1C) reflecting the absence of *Cst7* expression in the WT mouse brain in the absence of a pathological stimulus. Additionally, no tdTomato expression was detected in 5xFAD/ *Cst7*^DD-Cre^/Ai14^tdTomato^ mice not treated with TMP (Fig. 1D) supporting that there is no leakiness associated with the Cre line. Importantly, 5xFAD/ *Cst7*^DD-Cre^/Ai14^tdTomato^ mice treated with TMP display expression of tdTomato in plaque associated areas in both 4- month-old (Fig. 1E) and 11-month-old (Fig. 1F) cohorts, but not in areas outside the vicinity of plaques. In the younger cohort, tdTomato fluorescence colocalizes only with IBA1+ PAMs in the brain (Fig. 1G, H) and labels an average of 38% and 33% of PAMs in the subiculum and somatosensory cortex, respectively (Fig. 1I). In the older cohort, tdTomato fluorescence again colocalizes only with IBA+ PAMs (Fig. 1K), with a greater extent than in young mice, with an average of 78% and 58% of PAMs colocalizing with tdTomato in the subiculum and somatosensory cortex, respectively (Fig. 1L). The increase in the proportion of PAMs expressing tdTomato compared to the younger cohort, is consistent with the increase in DAM observed with disease progression in 5xFAD mice. To further characterize the tdTomato positive cells, we performed immunofluorescence (IF) staining for the DAM marker CD11c in the younger (Fig. 2A) and older cohorts (Fig. 2B) and found that an average of 84% and 86% of CD11c+ cells colocalized with tdTomato in the subiculum in young and aged cohorts, respectively (Fig. 2C), indicating our mouse line targets cells that express, or have expressed, DAM genes. We previously found that MAC2/LGALS3 stains both infiltrating myeloid cells in the brain, as well as a small subset of PAMs ^38^. Staining for MAC2 in young (Fig. 2D) and aged (Fig. 2E) cohorts shows that an average of 41% and 52% of MAC2+ cells colocalized with tdTomato in the subiculum in young and aged cohorts, respectively (Fig. 2F), indicating the existence of a distinct subset of MAC2^+^ PAMs that are not tdTomato^+^/CD11c^+^. Together, these results indicate that the novel *Cst7*^DD-Cre^ mouse line can effectively target inducible Cre expression within DAMs. The findings also suggest that the population of microglia that cluster around plaques may consist of functionally and transcriptionally different subsets of cells.

**Figure 2:**
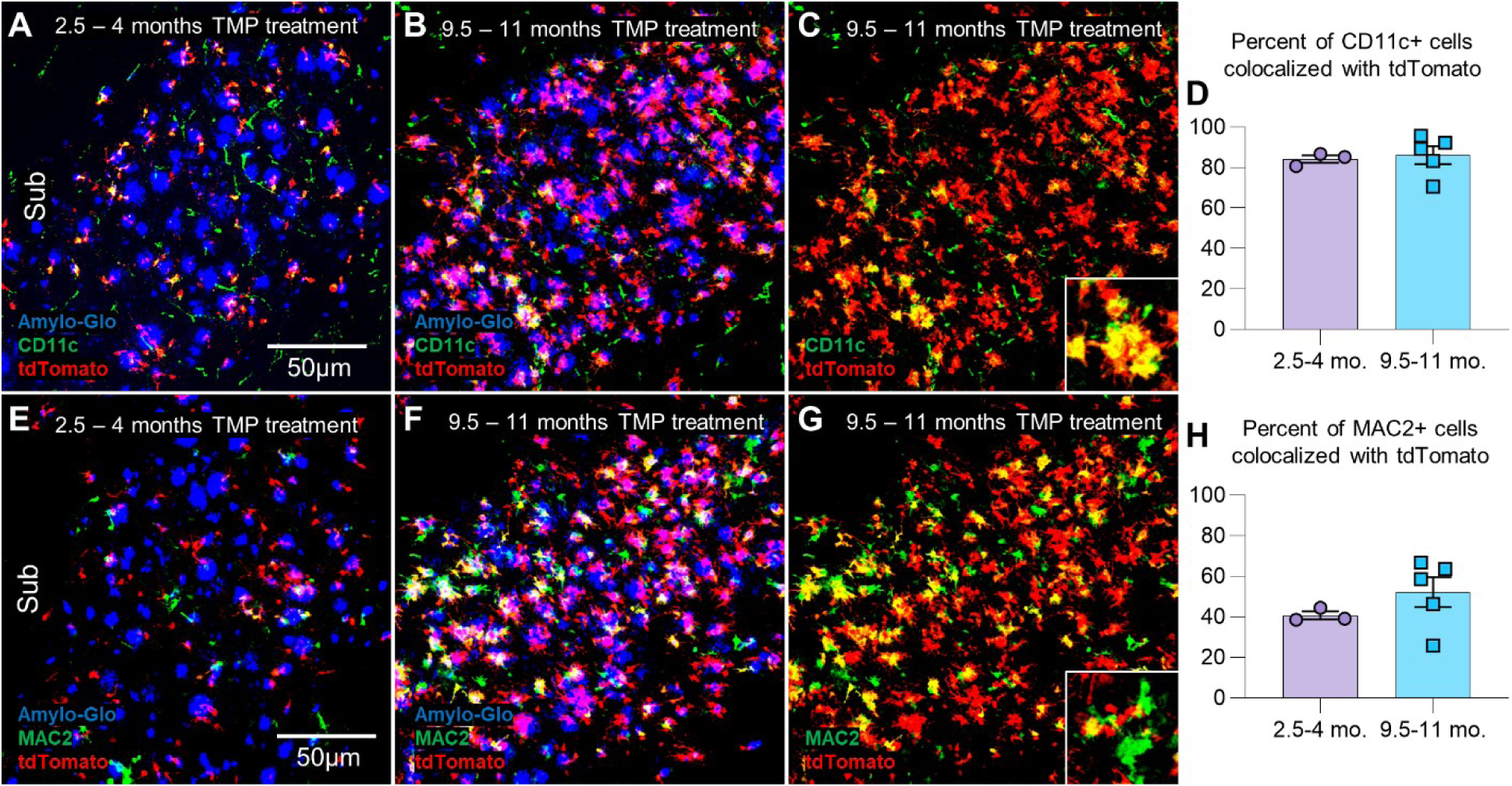
tdTomato+ cells colocalize more specifically to DAM markers. (A-D) Four-month-old (A) and 11–month-old (B, C) 5xFAD/*Cst7*^DD-Cre^/Ai14^tdTomato^ mice show colocalization of DAM marker CD11c with tdTomato. An average of 84% and 86% of CD11c+ cells are colocalized with tdTomato+ cells at 4 month and 11 months of age, respectively (D). (E-H) Four-month-old (E) and 11-month-old (F, G) 5xFAD/*Cst7*^DD-Cre^/Ai14^tdTomato^ mice show colocalization of infiltrating macrophage marker MAC2 with tdTomato. An average of 41% and 52% of MAC2+ cells are colocalized with tdTomato+ cells at 4 month and 11 months of age, respectively (H).

### *Cst7*^DD-Cre^ target DAMs associated with white matter in an inducible demyelinating cuprizone mouse model

While DAMs were first characterized in the 5xFAD mouse model ^3^, DAMs or DAM-like cells have since been found in other plaque models ^39,40^, tauopathy models ^40,41^, and other mouse models of neurodegeneration or demyelination^42–46^. We tested whether the *Cst7*^DD-Cre^ allele is capable of targeting DAMs in other disease-like states by utilizing cuprizone (CPZ), a copper-chelating chemical that causes oligodendrocytic cell death, demyelination, and brain wide neuroinflammation particularly in white-matter areas^47^. Ten-month-old *Cst7*^DD-Cre^/Ai14^tdTomato^ mice were treated with TMP in drinking water and a CPZ diet for 1.5 months (Fig. 3A). The brains of *Cst7*^DD-Cre^/Ai14^tdTomato^ mice treated with CPZ and TMP display cellular tdTomato fluorescence around white-matter areas in the brain (Fig. 3B), and all tdTomato expression colocalizes with IBA1+ microglia particularly in white-matter regions such as the striatum (Fig. 3C), and the corpus callosum (Fig. 3D). Similar to results using the 5xFAD transgene, tdTomato signal in CPZ-treated animals colocalizes with DAM marker CD11c (Fig. 3E) and infiltrating myeloid marker MAC2 (Fig. 3F). These findings support that our model can target DAMs in both the 5xFAD model and an inducible demyelinating mouse model.

**Figure 3:**
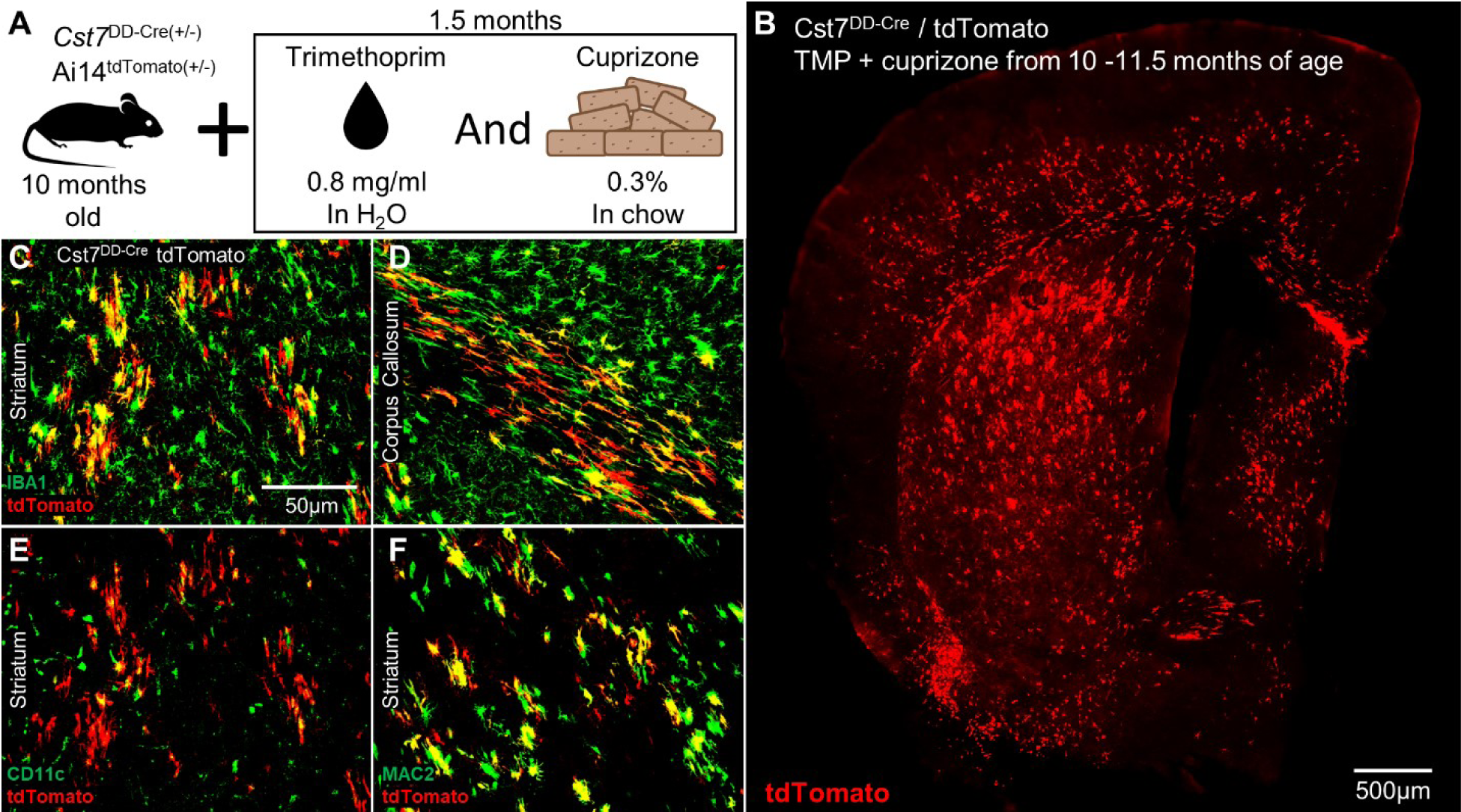
*Cst7*^DD-Cre^/Ai14^tdTomato^ mice fed a demyelinating cuprizone diet and treated with TMP show tdTomato expression in white matter-associated microglia. (A) Schematic showing experimental paradigm. Ten-month–old *Cst7*^DD-Cre^/Ai14^tdTomato^ mice were treated with TMP in drinking water and fed a cuprizone diet for 1.5 months before sacrifice. (B) Extensive tdTomato expression in white-matter areas of the brain in *Cst7*^DD-Cre^ / Ai14^tdTomato^ mice treated with CPZ and TMP. (C-D) Higher magnification images of the white matter areas of the brain, the striatum (C), and the corpus callosum (D), show colocalization of tdTomato with microglia around white matter. (E,F) Higher magnification images of the white matter areas of the striatum showing tdTomato colocalization with CD11c (E) and MAC2(F) markers of microglia.

### Spatial transcriptomics reveals tdTomato-positive microglia have higher DAM expression compared to other microglial sub-populations

In both TMP-treated 5xFAD *Cst7*^DD-Cre^/Ai14^tdTomato^ mice and *Cst7*^DD-Cre^/Ai14^tdTomato^ animals treated with TMP and CPZ, tdTomato expression was observed only in microglia demonstrating the specificity of the *Cst7-DD-Cre* allele to target this cell type. However, in both cases, not all microglia, including PAMs, expressed tdTomato. It is possible that not all microglia that expressed *Cst7* produced sufficient *Cst7*^DD-Cre^ activity to activate expression of the tdTomato lineage tracer. Alternatively, it is possible that only a sub-set of microglia expressed *Cst7* in these models. To discriminate between these and other possibilities we used spatial transcriptomics to characterize DAMs. To do so, we used Nanostring’s CosMx single cell spatial molecular imaging platform, utilizing a 1000 plex mouse neuroscience gene panel, plus custom probes for tdTomato. This method enables single-cell level gene expression analysis in all cell types, including microglia, overcoming limitations associated with single-cell and single-nucleus RNA-seq, such as technique-induced expression changes ^48–50^. Additionally, it can capture roughly 90% of cells within a given sample and provides information on genes at distinct spatial locations within the tissue, facilitating *in situ* counting and profiling of all cell populations within a brain slice. This method also enables analysis of gene expression in populations of microglia that may have expressed *Cst7* during their lifetime, but which might not currently be expressing *Cst7*. Analysis was performed on both the hippocampus and cortex of three (1F, 2M) 11-month-old 5xFAD/ *Cst7*^DD-Cre^/Ai14^tdTomato^, two (2M) 10-month-old CPZ-treated *Cst7*^DD-Cre^/Ai14^tdTomato^, and one (1M) 11-month-old *Cst7*^DD-Cre^/Ai14^tdTomato^ mice all treated with TMP (Fig. 4A). Cells are identified and segmented using histone, rRNA, GFAP, and DAPI (examples shown on Supp. Fig. S1A), then the transcripts of each gene per cell are measured.

**Figure 4:**
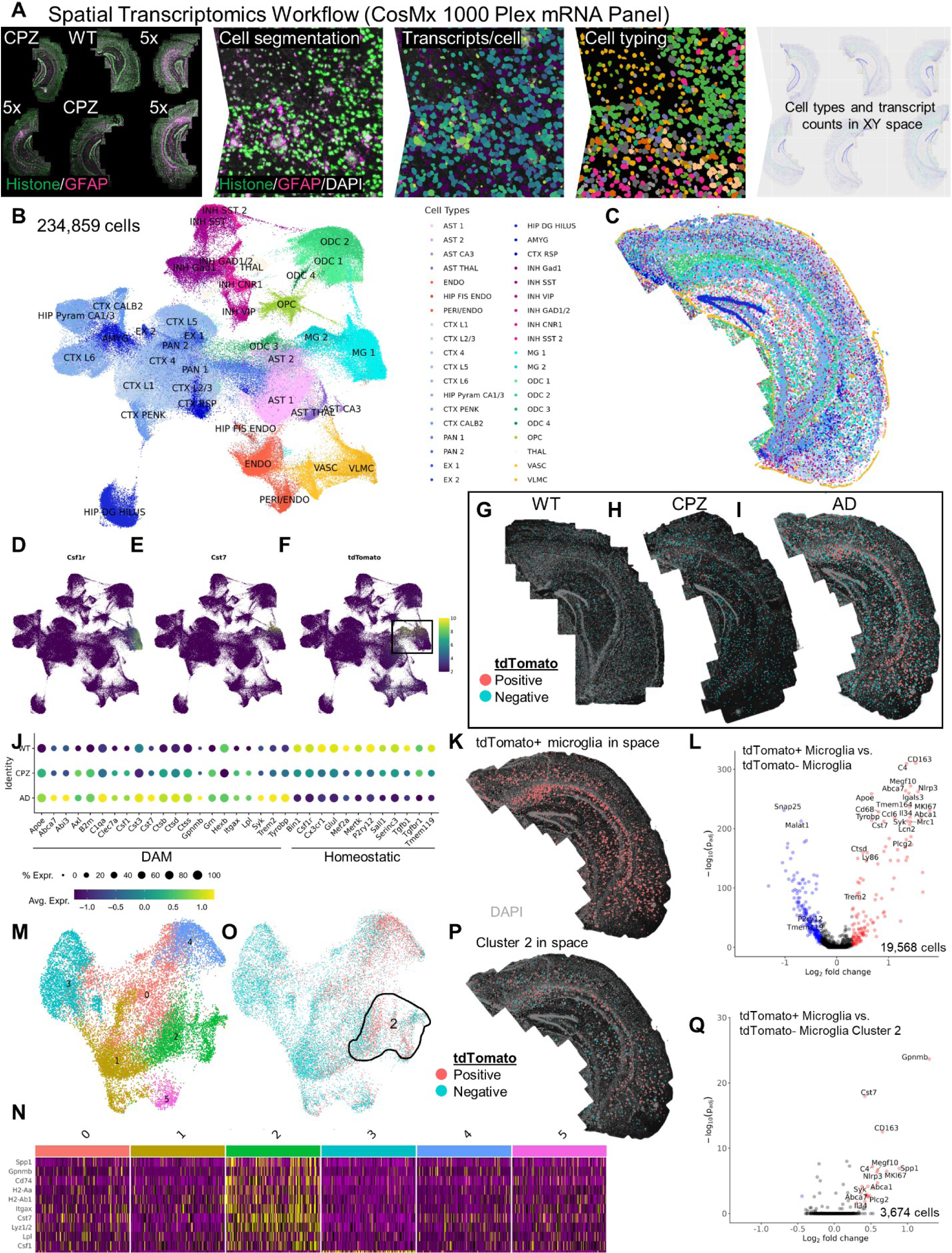
Spatial transcriptomic analyses reveal increased DAM gene expression in tdTomato^pos^ PAMs vs tdTomato^neg^ PAMs. (A) Schematic of spatial transcriptomics workflow using the CosMx 1000 Plex mRNA panel. Whole brains slices were mounted and fields of view (FOVs) of the hippocampus and cortex were selected. Cells were segmented using histone, GFAP, and DAPI and transcripts quantified in each cell for further analysis. (B-C) UMAP of 234,859 cells reveals 38 distinct clusters (B) that can be mapped in space on each brain (C). (D-F) UMAPs showing normalized expression of *Csf1r* (D), *Cst7* (E), and tdTomato (F), indicating these genes are expressed in MG 1 and MG 2, which are boxed in (F). (G-I) Clusters MG 1 and MG 2 mapped in space on a WT (G), cuprizone (CPZ), and 5xFAD brain with tdTomato^pos^ cells represented in red, and tdTomato^neg^ cells represented in cyan. (J) Dot Plot comparing microglial gene expression between WT, CPZ, and 5xFAD groups shows enriched DAM gene expression and diminished homeostatic microglial gene expression in CPZ and 5xFAD compared to WT. (K) tdTomato^pos^ cells from MG 1 and MG 2 mapped onto the 5xFAD brain. (L) Volcano plot comparing tdTomato ^pos^ microglia vs tdTomato^neg^ microglia from MG 1 and MG 2 shows increased DAM gene expression (e.g.: *Trem2*, *Cst7*, *Apoe*), and a decreased homeostatic gene expression (e.g.: *Tmem119*, *Pr2ry12*). (M-O) UMAP of 19,568 cells from MG 1 and MG 2 subclustered into six distinct clusters. (N) Heatmap showing the 10 most upregulated genes in cluster 2 compared to the other microglial clusters. DAM genes such as *Gpnmb*, *Cst7*, *Itgax*, and *Lpl* are highly upregulated in cluster 2 indicating this cluster may be associated with Aβ pathology. (O-P) UMAP showing tdTomato^pos^ and tdTomato^neg^ cells with cluster 2 encircled (O). tdTomato^pos^ and tdTomato^neg^ cells from cluster 2 mapped onto a 5xFAD brain shows that both types of cells are located in plaque associated areas shown in DAPI (P). (Q) Volcano plot comparing tdTomato^pos^ microglia vs tdTomato^neg^ microglia from cluster 2 shows an upregulation in DAM gene expression (e.g.: *Cst7*, *Gpnmb*, *Spp1*, *Itgax*).

To visualize the data, we first clustered a total of 234,859 cells using a shared nearest neighbor (SNN) modularity optimization-based clustering algorithm resulting in 38 distinct clusters and visualized these clusters using Uniform Manifold Approximation Projection (UMAP)(Fig. 4B) that were annotated according to gene expression and spatial location (Fig. 4C, Supp. Fig. S1B). Based on the expression of *Csf1r* (Fig. 4D), *Cst7* (Fig. 4E), and *tdTomato* (Fig. 4F) on the UMAP, we identified two clusters of microglia (MG 1 and MG 2) comprised of 19,568 cells. We then took the cells from MG 1 and MG 2, sorted them based on whether each cell expressed tdTomato, and mapped each cell onto *Cst7*^DD-Cre^/Ai14^tdTomato^ (Fig. 4G), *Cst7*^DD-Cre^/Ai14^tdTomato^ treated with CPZ (Fig. 4H), and 5xFAD/*Cst7*^DD-Cre^/Ai14^tdTomato^ (Fig. 4I) brains. As expected, very few tdTomato expressing (tdTomato^pos^) cells are seen in the *Cst7*^DD-Cre^/Ai14^tdTomato^ brain compared to the *Cst7*^DD-Cre^/Ai14^tdTomato^ CPZ-treated and 5xFAD/*Cst7*^DD-Cre^/Ai14^tdTomato^ brains, with the 5xFAD showing the most tdTomato^pos^ cells, suggesting that microglia in 5xFAD/*Cst7*^DD-Cre^/Ai14^tdTomato^ mice have the highest incidence of *Cst7* expression. These data are reinforced by the relative expression of DAM and homeostatic microglial genes, with *Cst7*^DD-Cre^/Ai14^tdTomato^ mice having the lowest expression of DAM genes (e.g. *Cst7*, *Apoe*, *Trem2*) and the highest expression of microglial homeostatic genes (e.g. *Tmem119*, *P2ry12*), while the 5xFAD/*Cst7*^DD-Cre^/Ai14^tdTomato^ mice have the highest DAM gene expression and lowest homeostatic microglial gene expression (Fig. 4J). We further characterized the difference between our tdTomato^pos^ DAM cells and all other microglia. As expected, the tdTomato^pos^ cells map onto areas of the brain where plaques are present (Fig. 4K). Differential gene expression (DGE) analysis of tdTomato^pos^ and tdTomato^neg^ microglia, shows that tdTomato^pos^ microglia have higher expression of DAM genes, and lower expression of homeostatic microglial genes, indicating that we can utilize *Cst7*^DD-Cre^ to target inflammatory microglial cells in the brain (Fig. 4L).

### tdTomato^pos^ microglia exhibit higher DAM gene expression compared to other PAMs

So far, we have shown that the tdTomato^pos^ cells in *Cst7*^DD-Cre^/Ai14^tdTomato^ mice have higher DAM gene expression compared to all other microglia. Next, we investigated whether these cells have higher DAM gene expression compared to other PAMs that do not express tdTomato, ignoring the homeostatic microglial population that was present in previous analysis. To do so, we further subclustered MG 1 and MG 2 into six distinct clusters (Fig. 4M). Of these subclusters, subcluster 2 had the highest expression of DAM genes (e.g. *Itgax*, *Cst7*, *Gpnmb*) compared to all other microglial subclusters (Fig. 4N). Subcluster 2 is also found to have a very high number of tdTomato^pos^ cells compared to other subclusters (Fig. 4O). Of note, there appears to be a high number of tdTomato^pos^ cells in subcluster 4 as well, and this subcluster is characterized by inflammation markers (e.g.: *CD163*, *Il34*, *Nlrp3*) (Supp. Fig. S2A). However, this subcluster appears to only be present in the 5xFAD/*Cst7*^DD-Cre^/Ai14^tdTomato^ group and is virtually absent in the *Cst7*^DD-Cre^/Ai14^tdTomato^ and *Cst7*^DD-Cre^/Ai14^tdTomato^ treated with CPZ groups (Supp. Fig. S2C). Upon viewing the cell segmentation of the 5xFAD brain stained for DAPI, it appears that subcluster 4 cells may be Aβ plaques themselves, which are stained by DAPI (Supp. Fig. S2D, E). Therefore, we decided to focus our analysis on subcluster 2. Upon mapping the 3,674 subcluster 2 cells that are either tdTomato^pos^ or tdTomato^neg^ onto the 5xFAD brain, we observe that these cells are specifically located in plaque associated areas (Fig. 4P). Importantly, DGE analysis of tdTomato^pos^ vs tdTomato^neg^ cells from this subcluster shows that tdTomato^pos^ cells have higher expression of DAM genes (e.g.: *Cst7*, *Gpnmb*, *Syk*, *Spp1*) (Fig. 4Q), indicating that the *Cst7*^DD-Cre^ can specifically be used to target DAMs while keeping other subsets of PAMs unaffected.

## Discussion

Genetic data show that microglia are both disease-driving and heterogeneous. Hence, it is important to understand the mechanisms of microglial heterogeneity and contributions of microglial subpopulations to better determine their role in AD. To that end, we aimed to develop a method to specifically target gene expression in DAMs while leaving other microglia and cell types unaffected. By doing so, we can better understand the specific contribution of these cells to AD, as the exact role of microglia in disease is currently unclear and there are confounds associated with targeting all microglia/myeloid cells as demonstrated by *Trem2* knock-out studies. From our own studies and single cell RNA sequencing studies, it is evident that microglia exhibit differential functions based on their RNA expression ^3,4,17–21,51^, and confusion about the role of microglia in AD may in part come from our inability to untangle the contribution of different microglial populations to AD.

We generated a novel mouse line with an allele of *Cst7* that expresses trimethoprim (TMP) inducible DD-Cre while retaining *Cst7* function. The *Cst7*^DD-Cre^ allele effectively allows Cre-dependent targeting of genetic function in DAMs, while sparing other microglia and cell types in the brains of 5xFAD mice. Our results suggest that DAM phenotypes are only induced next to plaques and these cells, and any progeny, do not migrate away from plaques after DAM expression, as has previously been hypothesized ^52^.

The *Cst7*^DD-Cre^ line provides distinct advantages over other traditional and inducible microglia-specific Cre models. Widely used microglia Cre lines such as *Cx3cr1*^CreER 53^, *Tmem119*^CreER 54^, and *P2ry12*^CreER 55^ lines have been invaluable in efforts to understand microglial biology. However, these lines target all microglial cells and tamoxifen can only be administered as a short-term treatment. Particularly, in the *Cx3cr1*^CreER^, the Cre-ER fusion protein disrupts CX3CR1 function and may have unintentional toxic effects that lead to microglial activation in mice ^53,56^. This line has also been shown to have Cre activity in the absence of tamoxifen ^57^. Recently, a novel *Clec7a*-CreER^T2^ mouse line was developed which can effectively label the DAM population, although tamoxifen is still the driver for Cre recombination at *loxP* sites, allowing only for short-term treatment ^58^. The *Cst7*^DD-Cre^ model avoids potential confounds of each of the other Cre lines, while specifically targeting DAMs with no Cre leakiness. *Cst7* encodes Cystatin F and is an endosomal/lysosomal cathepsin regulator and considered to be a DAM gene. Under normal conditions, CST7 is expressed by NK cells, T cells, and neutrophils in the periphery, with no expression found in the brain ^59^. In disease, CST7 is upregulated in microglia in white-matter during demyelination, and in microglia around plaques in amyloidosis models of disease ^3,39,59^. Additionally, CST7 expression in the brain appears to be dependent on TREM2 signaling as CST7 expression is abolished in TREM2 knockout mice ^3,41,60^. Although *Cst7* is highly upregulated in AD, little is known regarding its contribution to disease pathology. However, it has been shown that induction of its upstream kinase, RIPK1, increases CST7 levels and leads to impairment in lysosomal pathways.

In addition to showing enriched *Cst7* expression in PAMs, we observe an age-related increase in the proportion of tdTomato^pos^ PAMs at 11 months of age. Consistent with this, it has been shown that another DAM marker, CD11c, is elevated in 12-month-old 5xFAD mice compared to 4-month-old 5xFAD mice ^61^. This suggests that some microglia may require more time or exposure to Aβ to switch to a DAM gene profile, or that the microglial response to pathology is age-dependent, or possibly both, underlining the importance of studying the role of microglia at different stages of disease pathology. This is especially important at younger timepoints, as it appears there is more heterogeneity in PAMs at early timepoints when DAM expression is lower, versus later in disease when DAM expression appears higher. Additionally, the IHC and spatial transcriptomics data highlight the heterogeneity of PAMs themselves. Taken together, our data suggest that the dynamic expression of microglia at different disease stages and brain regions may have robust effects on therapeutic strategies targeting microglia, underlining the importance of studying the role of microglial subsets at different stages of disease pathology.

In summary, we have developed a novel mouse strain that represents a useful genetic tool to uncover the role of DAMs in AD and other diseases. The DD-Cre/TMP system is an innovative treatment approach with safer, long-term efficacy compared to traditional CreERT2/tamoxifen systems which may have detrimental effects on mouse safety and lead to increased activation of macrophages, all while only having the potential for short term treatment ^34,35^. Importantly, we have shown that DAMs represent a subset of PAMs and are characterized by higher levels of disease and inflammatory gene expression compared to other populations of microglia, including other PAMs.

## Acknowledgements

This work was supported by the National Institutes of Health (NIH) under awards: R01NS083801 (NINDS), R01 AG081599 (NIA), RF1AG056768 (NIA), and U54AG054349 (NIA Model Organism Development and Evaluation for Late-onset Alzheimer’s Disease [MODEL-AD]) to K.N.G, F31AG072852 (NIA) to C.M.H., T32NS082174 (NINDS) to K.I.T, and to T32NS121727 (NINDS) N.K. We acknowledge the support of the Chao Family Comprehensive Cancer Center Transgenic Mouse Facility Shared Resource, supported by the National Cancer Institute of the National Institutes of Health under award number P30CA062203. The content is solely the responsibility of the authors and does not necessarily represent the official views of the National Institutes of Health. Sources of funding did not have any role in study design, data collection or analysis, interpretation of results, or manuscript preparation or submission.

## Author Contributions

C.M.H. conceived and performed experiments, analyzed data, and wrote the manuscript. K.I.T. and N.K. performed experiments and analyzed data. S.K, V.S., G.R.M., J.N.N. and K.N.G. conceived experiments, provided supervision and wrote the manuscript. All authors read and approved the final manuscript.

