## Supplemental figure 1 and 2, and Supplemental file 1 for "Generation of an inducible destabilized-domain Cre mouse line to target disease associated microglia"

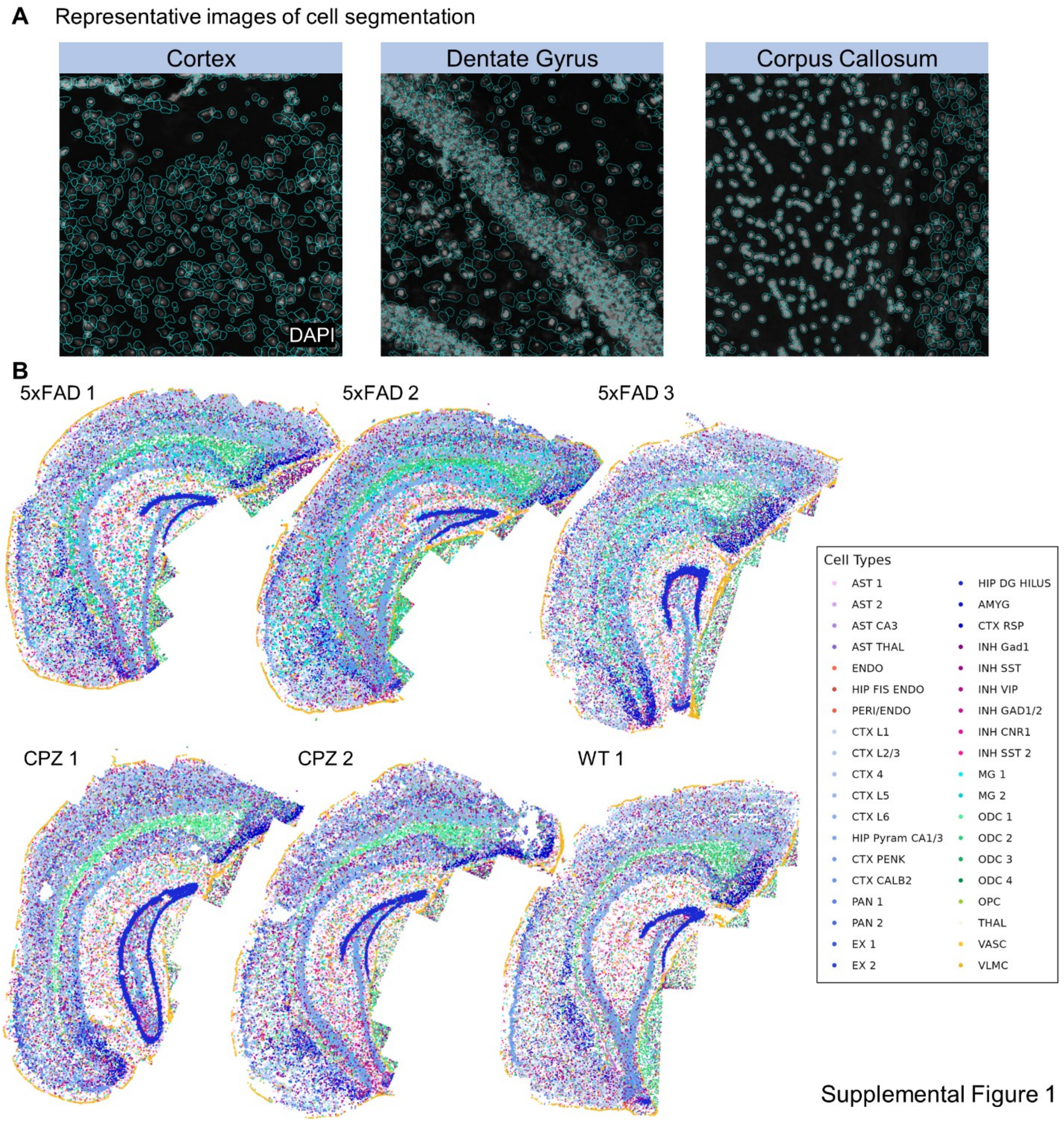


**Supplementary Fig. 1: CosMx cell segmentation and spatial mapping.**

(A) DAPI stained images showing representative cell segmentation in the cortex, dentate gyrus, and corpus callosum

(B) Cells from the the UMAP mapped onto each individual brain

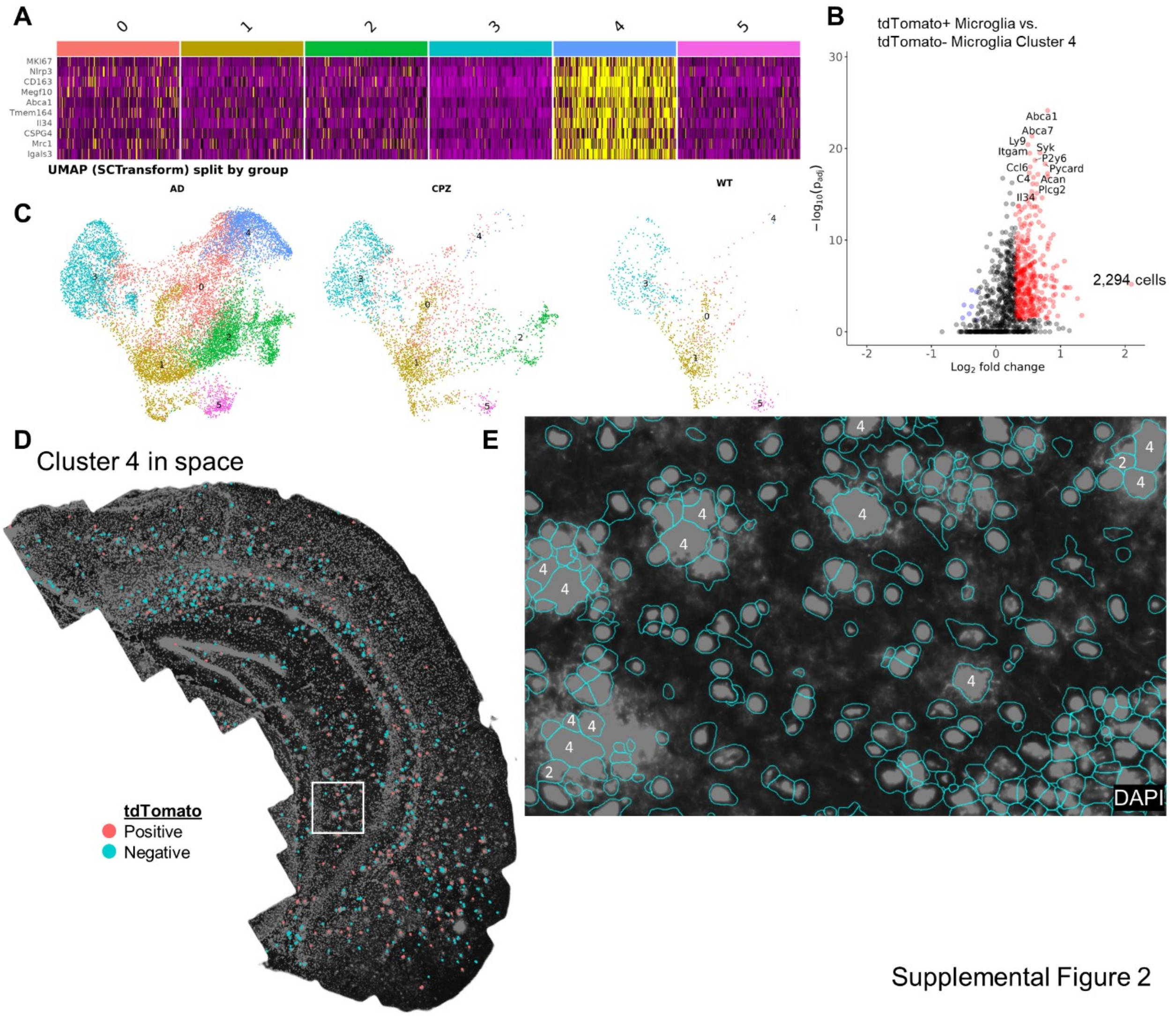


**Supplementary Fig. 2: Microglia subcluster 4 is comprised of Aβ plaques.**

(A) Heatmap showing 10 must upregulated genes in in cluster 4 compared to the other microglial clusters.

(B) Volcano plot comparing tdTomato$^{pos}$ microglia vs tdTomato$^{neg}$ microglia from cluster 2 shows increased inflammatory gene expression.

(C) UMAP split by 5xFAD, CPZ, and WT groups showing that cluster 4 is predominantly present in the 5xFAD group.

(D-E) Microglial subset cluster 4 tdTomato$^{pos}$ and tdTomato$^{neg}$ cells mapped onto a 5xFAD brain (D). White box indicates area of zoomed in FOV shown in (E).

(E) Cell segmentation image of a 5xFAD brain showing that some plaques stain positive by DAPI and are counted as cells which are subsequently grouped into microglial subset cluster 4.

Supplementary File 1

Sequence of 1805 base ssODN used to insert DD-Cre-P2A coding sequence into *Cst7* exon 1

ecDHFR DD linker iCRE P2A- exon 1

100 bp homology arms

AGCAAGAAACCACGCCCCACTGAAGCTACCCCACCATGCCCTGGTCCTGGAGCTGTACTTGCCGAGCACT
TGGCCCACTACATGCTCCCTGCATTTCCCCAGCCATGATCTCTCTGATTGCCGCTCTGGCCGTGGACTAC
GTGATCGGGATGGAAAACGCTATGCCATGGAATCTGCCCGCCGATCTGGCTTGGTTCAAGAGGAACACCC
TGAACAAGCCAGTGATCATGGGCAGACACACTTGGGAGTCCATTGGCCGGCCCCTGCCTGGACGCAAGAA
CATCATTCTGAGCTCCCAGCCCTCTACCGACGACAGGGTGACATGGGTGAAAAGTGTGGACGAAGCCATT
GCCGCTTGCGGAGATGTGCCCGAGATCATGGTCATCGGCGGAGGGAGAGTGATCGAGCAGTTCCTGCCTA
AGGCCCAGAAACTGTACCTGACTCACATTGACGCTGAGGTGGAAGGGGACACCCATTTTCCTGATTATGA
GCCAGACGATTGGGAAAGCGTGTTCTCCGAGTTTCACGACGCCGATGCTCAGAATTCTCATAGTTATTGC
TTTGAGATCCTGGAAAGGAGAGGCGCGCCTAAGAAGAAGAGGAAAGTCTCCAACCTGCTGACTGTGCACC
AAAACCTGCCTGCCCTCCCTGTGGATGCCACCTCTGATGAAGTCAGGAAGAACCTGATGGACATGTTCAG
GGACAGGCAGGCCTTCTCTGAACACACCTGGAAGATGCTCCTGTCTGTGTGCAGATCCTGGGCTGCCTGG
TGCAAGCTGAACAACAGGAAATGGTTCCCTGCTGAACCTGAGGATGTGAGGGACTACCTCCTGTACCTGC
AAGCCAGAGGCCTGGCTGTGAAGACCATCCAACAGCACCTGGGCCAGCTCAACATGCTGCACAGGAGATC
TGGCCTGCCTCGCCCTTCTGACTCCAATGCTGTGTCCCTGGTGATGAGGAGAATCAGAAAGGAGAATGTG
GATGCTGGGGAGAGAGCCAAGCAGGCCCTGGCCTTTGAACGCACTGACTTTGACCAAGTCAGATCCCTGA
TGGAGAACTCTGACAGATGCCAGGACATCAGGAACCTGGCCTTCCTGGGCATTGCCTACAACACCCTGCT
GCGCATTGCCGAAATTGCCAGAATCAGAGTGAAGGACATCTCCCGCACCGATGGTGGGAGAATGCTGATC
CACATTGGCAGGACCAAGACCCTGGTGTCCACAGCTGGTGTGGAGAAGGCCCTGTCCCTGGGGGTTACCA
AGCTGGTGGAGAGATGGATCTCTGTGTCTGGTGTGGCTGATGACCCCAACAACTACCTGTTCTGCCGGGT
CAGAAAGAATGGTGTGGCTGCCCCTTCTGCCACCTCCCAACTGTCCACCCGGGCCCTGGAAGGGATCTTT
GAGGCCACCCACCGCCTGATCTATGGTGCCAAGGATGACTCTGGGCAGAGATACCTGGCCTGGTCTGGCC
ACTCTGCCAGAGTGGGTGCTGCCAGGGACATGGCCAGGGCTGGTGTGTCCATCCCTGAAATCATGCAGGC
TGGTGGCTGGACCAATGTGAACATtGTGATGAACTACATCAGAAACCTGGACTCTGAGACTGGGGCCATG
GTGAGGCTGCTCGAGGATGGGGACTGAGGAAGCGGAGCTACTAACTTCAGCCTGCTGAAGCAGGCTGGAG
ACGTGGAGGAGAACCCTGGACCTTGGCTcGCtATcCTGCTTGCCCTCTGCTGCCTAACTTCTGACACCCA
TGGGGCACGCCCCCCAGGTAAGAGAAGGGTCATCTGGCCCCATCCCAGAGTACTG

**Supplementary File 1:** Annotated sequence of 1805 base ssODN used to insert DD-Cre-P2A coding sequence into *Cst7* exon 1

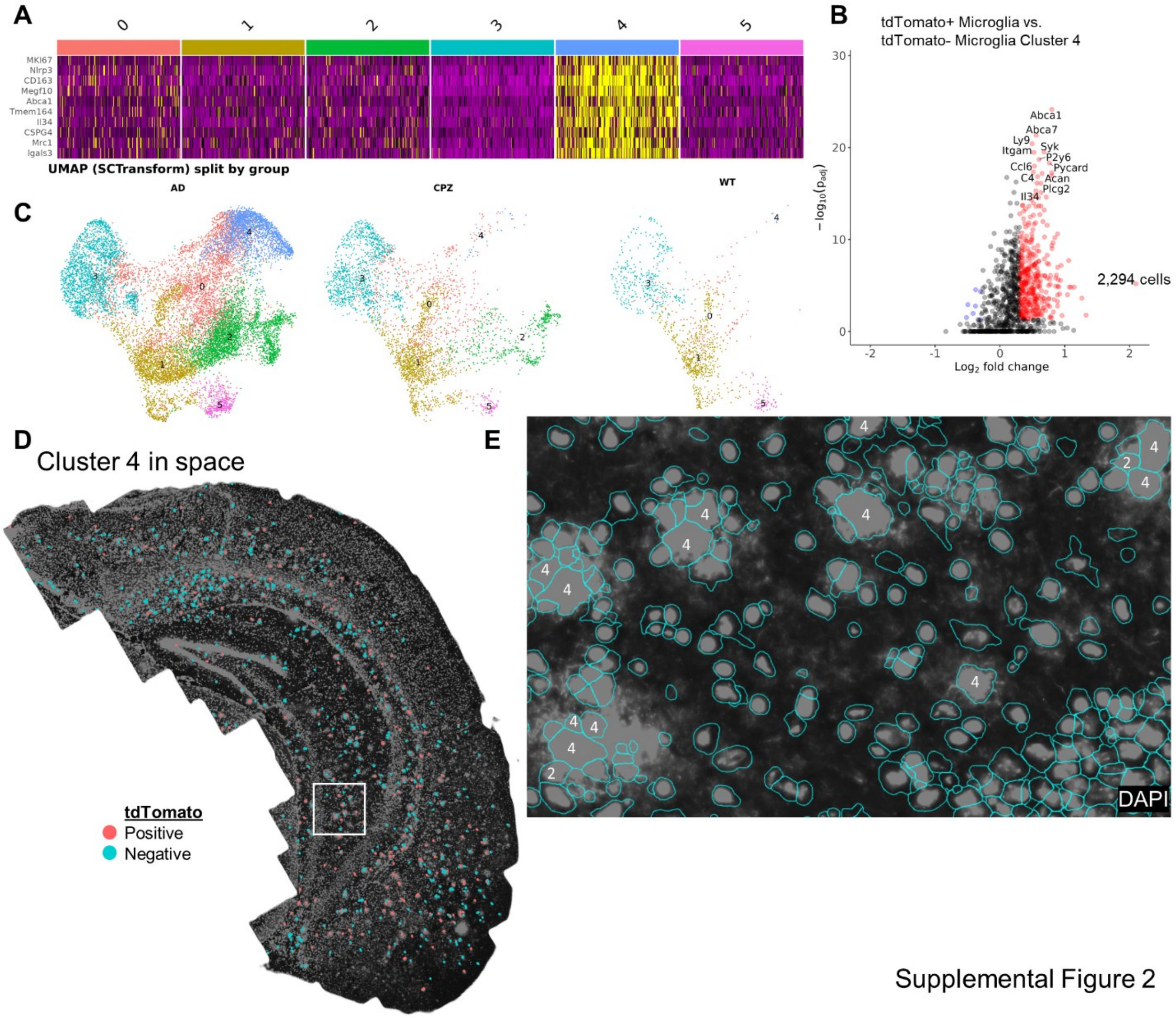


Supplemental Figure 2
